# Dysregulation of Long Non-coding RNA (lncRNA) Genes and Predicted lncRNA-Protein Interactions during Zika Virus Infection

**DOI:** 10.1101/061788

**Authors:** Arunachalam Ramaiah, Deisy Contreras, Vineela Gangalapudi, Masumi Sameer Padhye, Jie Tang, Vaithilingaraja Arumugaswami

## Abstract

Zika Virus (ZIKV) is a causative agent for poor pregnancy outcome and fetal developmental abnormalities, including microcephaly and eye defects. As a result, ZIKV is now a confirmed teratogen. Understanding host-pathogen interactions, specifically cellular perturbations caused by ZIKV, can provide novel therapeutic targets. In order to complete viral replication, viral pathogens control the host cellular machineries and regulate various factors, including long noncoding RNA (lncRNA) genes, at transcriptional levels. The role of lncRNA genes in the pathogenesis of ZIKV-mediated microcephaly and eye defects is currently unknown. To gain additional insights, we focused on profiling the differentially expressed lncRNA genes during ZIKV infection in mammalian cells. For this study, we employed a contemporary clinical Zika viral isolate, PRVABC59, of Asian genotype. We utilized an unbiased RNA sequencing approach to profile the lncRNA transcriptome in ZIKV infected Vero cells. We identified a total of 121 lncRNA genes that are differentially regulated at 48 hours post-infection. The majority of these genes are independently validated by reverse-transcription qPCR. A notable observation was that the lncRNAs, MALAT1 (Metastasis Associated Lung Adenocarcinoma Transcript 1) and NEAT1 (Nuclear Paraspeckle Assembly Transcript 1), are down-regulated upon Zika viral infection. MALAT1 and NEAT1 are known as nuclear localized RNAs that regulate gene expression and cell proliferation. Protein-lncRNA interaction maps revealed that MALAT1 and NEAT1 share common interacting partners and form a larger network comprising of 71 cellular factors. ZIKV-mediated dysregulation of these two regulatory lncRNAs can alter the expression of respective target genes and associated biological functions, an important one being cell division. In conclusion, this investigation is the first to provide insight into the biological connection of lncRNAs and ZIKV which can be further explored for developing antiviral therapy and understanding fetal developmental processes.

## INTRODUCTION

The human mosquito-borne pathogen, Zika Virus (ZIKV), is strongly associated with fetal developmental abnormalities, such as microcephaly and eye defects, as well as poor pregnancy outcomes [1–3]. The teratogenic ZIKV belongs to *Flaviviridae* family and the viral genome comprises of a positive-sense, single-stranded RNA molecule that encodes a large polyprotein. This polyprotein is proteolytically cleaved into 11 functional peptides of viral structural and non-structural components. ZIKV was first isolated from a Rhesus monkey in 1947 [4, 5]. The recent adaptation of ZIKV to a human host resulted in several epidemics across the globe [6, 7]. The virus is transmitted by mosquitos and through sexual contact. The major concern is the increasing geographical distribution of mosquito vectors *Aedes aegypti*, and *Aedes albopictus*, which is responsible for ZIKV transmission in humans. Understanding the interaction of ZIKV with its human host is crucial for developing effective vaccines and therapeutic measures.

The main focus of this study is to discern the effect of Zika viral infection on the expression of long non-coding RNA (lncRNA) genes. The presence of lncRNAs has been frequently determined with high confidence from transcriptomes and microarray technologies [8]. The Encyclopedia of DNA Elements (ENCODE) Project Consortium (GENCODE release 24) estimates that the human genome incorporates around 62% of 25,823 ncRNA genes as lncRNA genes that encode 28,031 well-defined lncRNA transcripts (<u>http://www.gencodegenes.org</u>). The minority of the non-coding transcripts are small, whereas, the majority of them exceed 200 nucleotides in length, and they are consequently cataloged as lncRNAs [9]. While this practical definition is fairly arbitrary, the size cutoff clearly differentiates lncRNAs from small regulatory RNAs, such as microRNAs (miRNAs), endogenous small interfering RNAs (endo-siRNAs) that participate in RNA interference (RNAi), and Piwi-associated RNAs (piRNAs). It is noted that the classical ncRNAs are ribosomal RNAs (rRNAs), ribozymes, transfer RNAs (tRNAs), small nuclear RNAs (snRNAs), small nucleolar RNAs (snoRNAs), and telomere-associated RNAs (TERC, TERRA) [10]. The majority of the lncRNA transcripts are expressed from intergenic regions and, to a lesser extent, in the protein-coding gene regions as sense or antisense transcripts.

Previous studies have provided an unbiased finding of non-coding transcripts across many cell types and tissues [11, 12] and made the conclusion that a vast majority of the genome was transcribed [13]. Interestingly, lncRNAs are playing a critical role in the developmental processes and etiology of a wide range of diseases. The expression patterns of many lncRNAs are associated with numerous key cellular processes, such as embryonic stem cell pluripotency, immune response, regulation of the cell cycle, and diseases such as cancer [9, 14–18]. However, the function of the majority of lncRNAs remains unknown [10]. Even though lncRNAs exert a spectrum of regulatory mechanisms across different cellular pathways, a broad theme is emerging in which lncRNAs are driving the assembly of RNA-protein complexes that can facilitate the regulation of gene expression. Knowing that there is an interaction between lncRNAs and protein coding genes in both normal and disease states is essential to uncover the key functional roles of lncRNAs in cellular and developmental processes [18, 19]. Thus, we carried out transcriptome analysis to determine expression patterns of lncRNAs in the ZIKV infected Vero cells that serve as a mammalian cell model. This study provides new insight into the interaction between ZIKV and its host.

## MATERIALS AND METHODS

### Cells

The Vero cell line (ATCC, USA) (kidney cells from an African green monkey) was cultured using complete Dulbecco’s modified Eagle’s medium (DMEM) (Fisher Scientific, USA). DMEM was supplemented with 10% fetal bovine serum (FBS), penicillin (100 units/ml), streptomycin (100 mg/ml), 10 mM Hepes, and 2 mM L-glutamine (Fisher Scientific, USA). The cells were cultured at 37°C with 5% CO_2_ conditions and passaged every second or third day for experimental assays or when the cells reached over 80% confluency.

### Zika virus

A ZIKV clinical isolate, PRVABC59 (GenBank accession number KU501215), belonging to an Asian genotype was used in this study [20]. Early passage virus was obtained from the Centers for Disease Control and Prevention (CDC). Viral stocks were generated in Vero cells. Viral titer was estimated by a plaque assay, as reported previously [21].

### Zika viral infection

Vero cells were seeded in a 12-well plate at a density of 1 × 10^5^ cells per well. The following day, Zika viral inoculum was prepared in serum-free DMEM media at a multiplicity of infection (MOI) of 1 and was added to each well in triplicate. To allow virus attachment and entry, the plates were incubated in 37°C with 5% CO_2_ for 4-6 hrs. After the designated incubation period, the inoculum was replaced with serum supplemented media. At 12, 24 and 48 hrs postinoculation, cells were harvested for RNA isolation. Uninfected control (mock) cells were included for each time point for comparison and were routinely treated with such the media without any viral inoculum.

### RNA sequencing sample preparation

Total RNA was isolated from the cells using an RNeasy Mini Kit (QIAGEN, USA). RNase-Free DNase treatment was performed on the column to remove residual DNA. Fourteen RNA samples [duplicate mock and infected samples for 12 and 24 hr time points (total 8 samples) and triplicate samples for 48 hr time point (6 samples)] were assessed for concentration and quality using the Thermo Scientific (Waltham, MA) NanoDrop 8000 Spectrophotometer. RNA sequencing libraries were constructed using Illumina TruSeq RNA Sample Preparation Kit v2 (Illumina, San Diego, CA) per manufacturer’s instructions. Briefly, total RNA with RNA integrity (RIN) scores of 9 or better using Agilent Bioanalyzer RNA 6000 Nano kit were used for RNA library preparation. The poly (A)+ RNA was then purified from one microgram of total RNA from each sample using oligo-dT attached magnetic beads with two rounds of purification. Subsequently, the poly (A)+ RNA was fragmented and primed for cDNA synthesis according to manufacturer’s recommendations. RNA adapters and barcodes were ligated to cDNA to allow for clonal amplification and multiplexing. Sequencing was done on an Illumina NextSeq 500 using 75bp single-end sequencing kit to yield an average read depth of 28 million reads per sample with a minimum number of reads for a sample of at least 16 million reads. The sequencing data were deposited to Gene Expression Omnibus (GEO) database with the accession number GSE83900.

### Data Analysis of RNA sequencing

Raw reads obtained from RNA-seq were aligned to the human genome using STAR (version 2.5) [22] with a custom human GRCh38 transcriptome reference downloaded from http://www.gencodegenes.org, which contains all protein-coding and long non-coding RNA genes based on human GENCODE version 23 annotation. A less stringent set of mapping parameters (allowing 5 mismatches per 75 bp reads) was used to align primate Vero cells to human reference.

Expression counts for each gene in all of the samples were normalized by a modified trimmed mean of the M-values normalization method. The unsupervised principal component analysis (PCA) was performed with DESeq2 Bioconductor package version 1.10.1 in R version 3.2.2 [23]. Each gene was fitted into a negative binomial generalized linear model. The Wald test was applied to assess the differential expressions between two sample groups by DESeq2. The Benjamini and Hochberg procedure [24] was applied to adjust for multiple hypothesis testing and differential expressed gene candidates were selected with a false discovery rate of less than 0.10. For visualization of coordinated gene expression in the samples, a hierarchical clustering with the Pearson correlation distance matrix was performed with differential expressed gene candidates using the Bioconductor gplots package (version 2.14.2) in R. Significantly expressed genes were assessed for pathway enrichment using DAVID release 6.7 (https://david.ncifcrf.gov/) and Ingenuity Pathway Analysis. (http://www.ingenuity.com/products/ipa) (QIAGEN, Redwood City). The significantly enriched canonical pathways were defined as having a q-value of <.01.

### Reverse transcription-quantitative PCR (RT-qPCR) Analysis

Total RNA was extracted from uninfected and ZIKV-infected cells using an RNeasy Mini Kit (QIAGEN). cDNA was synthesized using 3μg of total RNA and SuperScript III Reverse Transcriptase kit (Fisher Scientific, USA) and random hexamer primers as recommended by the manufacturers. Gene expression was quantified using Platinum SYBR Green qPCR SuperMix-UDG with ROX Kit (Life Technologies) by QuantStudio^TM^ 12K Flex Real-Time PCR System (Life Technologies). The reaction for each gene expression was completed in duplicate and housekeeping human gene, Glyceraldehyde 3-phosphate dehydrogenase (GAPDH), was used as an internal reference control. The list of qPCR primers and their sequences are given in Supplementary Table 1. The following conditions were used for real-time amplification of the cDNA: 50°C for 2 min; 95°C for 2 min followed by 40 cycles of 95°C for 15 sec and 60°C for 1 min. The mRNA expression of each target gene was calculated based on 2^−ΔCt^ values and fold-change was measured to the endogenous control, GAPDH.

### Immunofluorescence Assay

Both the mock and infected cells were fixed with methanol for 30 mins in −20°C. Following three PBS washes, the cells were blocked with a blocking buffer (10% fetal bovine serum, 3% BSA, 0. 1% Triton-x 100 in PBS). Subsequently, the cells were incubated with mouse monoclonal antibody for Flavivirus group antigen [D1-4G2-4-15 (4G2)] (Absolute Antibody Ltd.) at a 1:200 dilution overnight at 4°C. The secondary antibody, mouse anti-goat polyclonal antibody-FITC (Santa Cruz Biotechnology) (Life Technologies, USA) was added at a 1:1000 dilution and was incubated for 1 hr at room temperature. Between antibody changes, the cells were washed five times with PBS. The nuclei were stained with a DAPI dye (Life Technologies, USA).

### Non-coding RNA and protein interaction networks

To identify the high confident lncRNA-protein interaction network for NEAT1 and MALAT1, we used RNA-protein Association and Interaction Network’s (RAIN) web interface (http://rth.dk/resources/rain/). Subsequently, the combined NEAT1 and MALAT1 interaction analyses were carried out in the Search Tool for the Retrieval of Interacting Genes/Proteins (STRING) v10.0 database [25]. To avoid false positive results, a high confidence interaction score of 0.7 was used as a cut-off to obtain a maximum of 20 interactions under *evidence mode* for each of the two lncRNAs. The combined interacting network of these two lncRNAs with other human proteins was also predicted. The comprehensive interaction network was predicted using the 11 identified differentially co-expressed genes in normal and ZIKV infected conditions. Typically, the interaction network view summarizes the predicted associations for a specific group of proteins. The network nodes are proteins whereas the edges represent the predicted functional associations. The *evidence mode* based edges are drawn in with up to 7 differently colored lines representing the existence of the seven types of evidences. The evidence falls under 3 broad categories, namely known interactions, predicted interactions and other interactions, which was used in predicting the associations. The known interactions include the evidence that was experimentally determined (pink) or the evidence that was curated from database (sky blue). The predicted interactions depict the evidence of gene neighborhood (green), gene fusion (red) or gene co-occurrence (blue). The other interactions show evidence of co-expression (black), text-mining (light green) and protein homology (purple). Larger nodes reflect the availability of structural information associated with the factors. The Kyoto Encyclopedia of Genes and Genomes (KEGG) pathway analysis was done based on factors in the interaction networks.

### Statistical analysis

Error bars in the graph reflect the standard deviation. P-values were determined by the twotailed Student’s t-test and significance was reported if the p value is <0.05.

## RESULTS

### Zika virus infectious cell culture system

We utilized a Vero cell-based *in vitro* infectious culture system to investigate the ZIKV effect on the expression of lncRNAs. Vero cells supported robust lytic infection by the Asian ZIKV genotype (Fig. 1A). The viral-mediated cytopathic effects (CPEs), consisting of cell rounding and lysis, were observed at 48 hpi. The Zika viral genome replication was quantified by RT-qPCR, as described in the Methods section. At 48 hpi, several log increases in the ZIKV genome replication was observed (Fig. 1B). The virus infection in the Vero cells was also confirmed by detecting viral envelope proteins in the cytoplasm by Immunofluorescence Assay (Fig. 1C). These virological assays confirm the *in vitro* establishment of Zika virus infection, as well as a visible impact on cell health. Subsequently, we investigated the ZIKV-specific alterations in cellular gene expression profiles.

**Figure 1.**
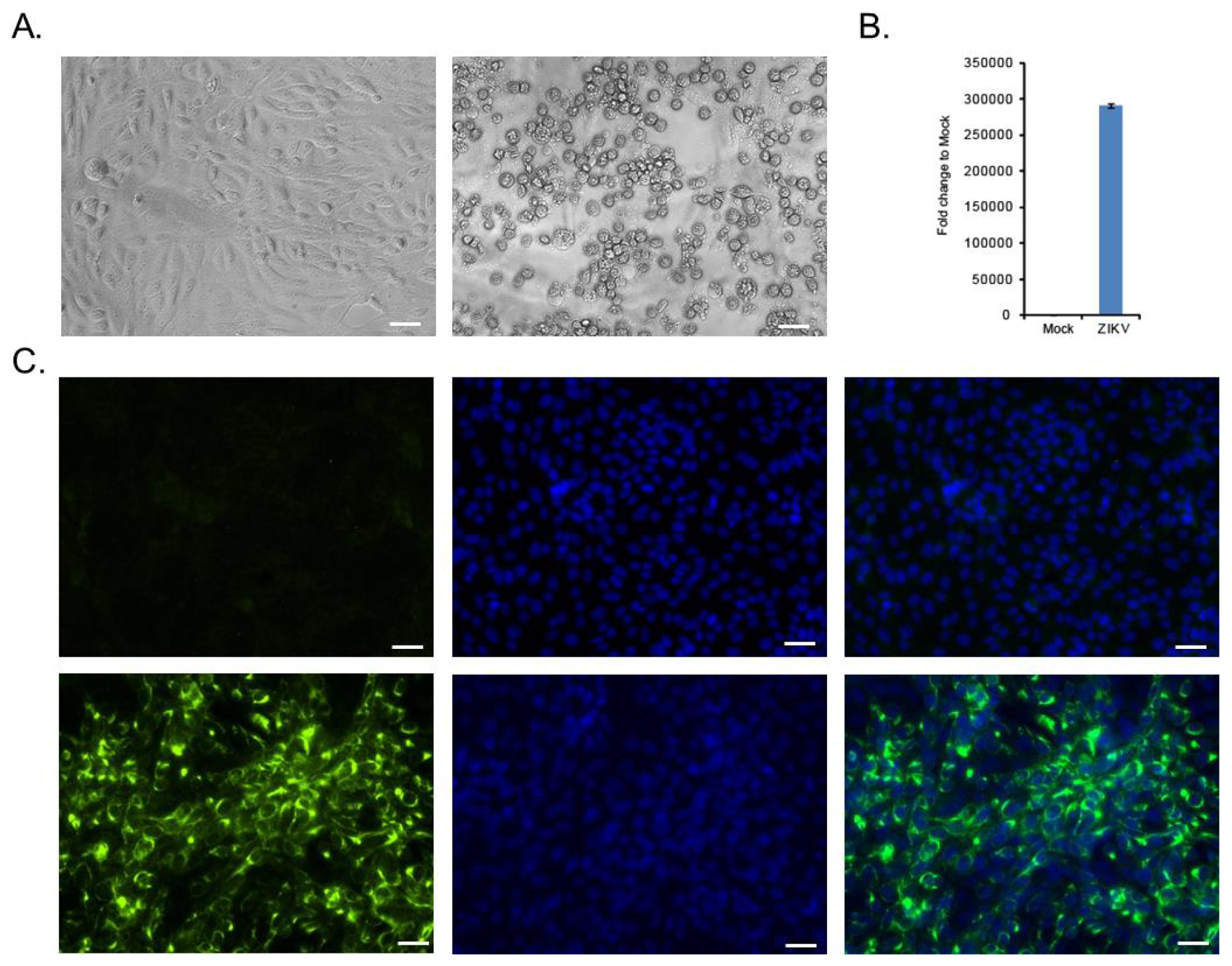
Vero cell-based ZIKV infectious culture system. (A) Bright field image presents ZIKV infected cells at 48 hours post-infection (hpi). Uninfected control (mock) cells are shown as well. (B) Graph depicts Zika viral genome replication at 48 hpi, and fold-change was calculated to that of mock control cells. (C) Immunofluorescence analysis shows ZIKV infected cells are positive for viral envelope protein.

### Transcriptome analysis of Zika viral infected cells

The transcriptome of all 14 samples from the Vero cells both with and without Zika viral infection were characterized at three different time points (12, 24, and 48 hpi) by RNA-Seq. After mapping with human transcriptome, 65-69% of overall mapping was observed per sample, >80% of the reads were successfully mapped to the exonic region in each sample and >80% of the aligned reads were uniquely mapped, which confirms the high quality of our samples and sequencing. We focused on identifying lncRNA mRNA that are differentially expressed between uninfected and ZIKV infected cells. The changes in lncRNA expression were observed only at the later time point of 48 hpi. For the 48 hpi time point, a total of 121 lncRNA genes were differentially expressed with an adjusted p value of <0.05 and 87 of these genes had an adjusted p value of <0.01 (Supplementary Table 2). A heat map of these differentially expressed lncRNAs is given in Figure 2.

**Figure 2.**
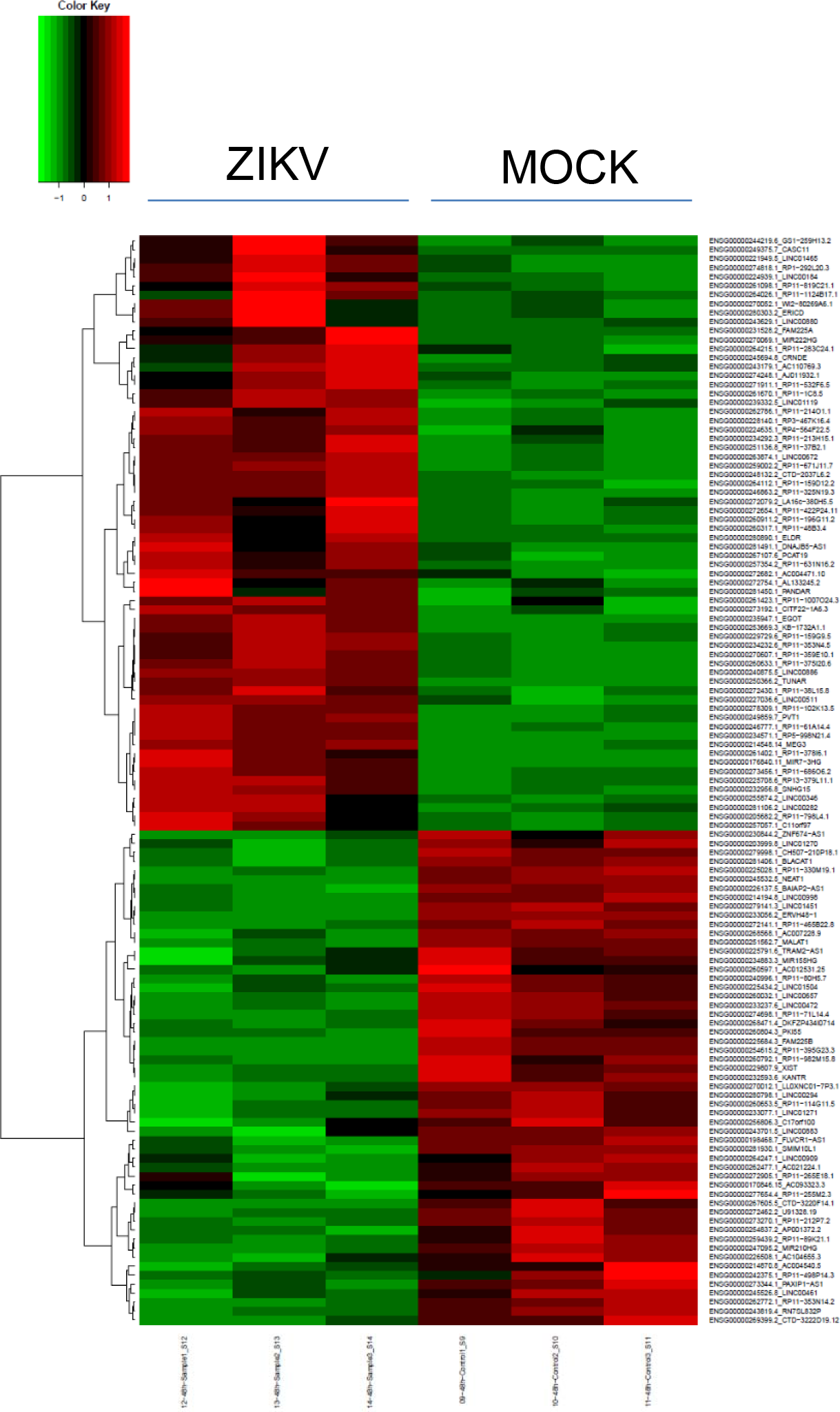
Global transcriptional analysis of ZIKV infected cells. Heat map depicts the differentially expressed lncRNA genes at 48 hour post-infection. (Red: up-regulated genes; Green: down-regulated genes). The hierarchical clustering was done using 121 DE lncRNAs for infected and control samples.

### Validation of lncRNAs differentially expressed during Zika virus infection

For validation, ZIKV infection experiment was repeated in Vero cells. Total RNA isolated at 48 hpi was subjected to RT-qPCR analysis to verify the lncRNAs. We confirmed the following up-regulated genes: LINC00184, LINC00672, RP11-213H15.1, EGOT, RP11-671J11.7, MIR222HG, KB-1732A1.1, C11orf97 and CITF22-1A6.3 (Fig. 3). Several of the down-regulated genes were also verified. Notably, MALAT1 (Metastasis Associated Lung Adenocarcinoma Transcript 1) and NEAT1 (Nuclear Paraspeckle Assembly Transcript 1) lncRNA levels were highly reduced in Zika virus infected cells (Fig. 3). HUCL (Hepatocellular Carcinoma Up-Regulated Long Non-Coding RNA) lncRNA was included as a control, which did not show changes during ZIKV infection. Since most of the lncRNAs are functionally uncharacterized, we focused on the previously known NEAT1 and MALAT1 for downstream studies.

**Figure 3.**
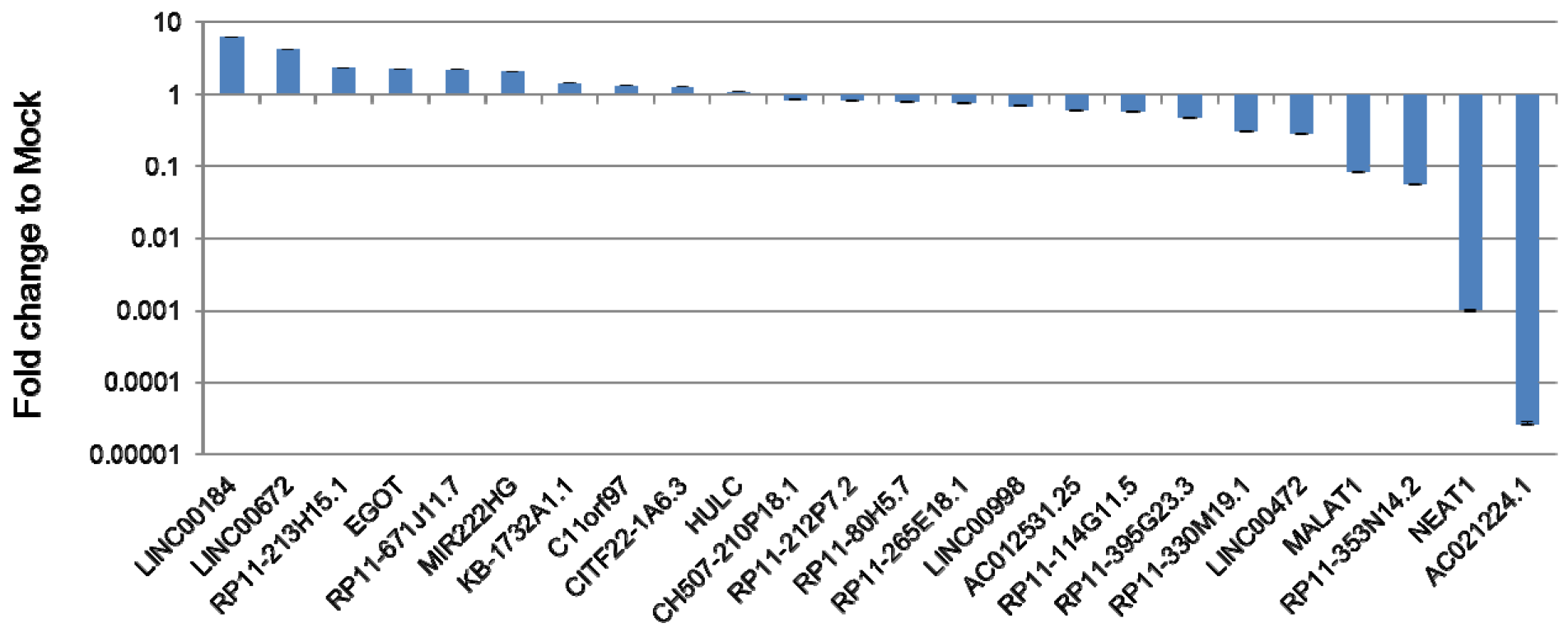
Validation of differentially expressed lncRNAs. Graph shows the expression levels of selected lncRNAs in ZIKV infected cells. Gene expression was quantified by RT-qPCR. Housekeeping gene GAPDH was used to normalize mRNA expression between ZIKV infected and mock control cells. For each gene, fold-change was calculated to that of uninfected cell control (mock) and presented as a bar graph.

### Functional interactions of differentially expressed lncRNAs and proteins in ZIKV infected cells

Identifying an interaction network of lncRNAs and protein coding genes, which are differentially regulated in ZIKV infected cells relative to the uninfected control, is essential to uncover the key biological functions of lncRNAs in cell injury processes. Hence, we focused on identifying the factors that can interact with lncRNAs NEAT1 and MALAT1 using RAIN and STRING. Initial seed interaction networks for NEAT1 and MALAT1 predicted a maximum of 20 interacting factors (Supplementary Figures 1 and 2). There were 20 and 19 high confidence nodes that involved in 31 and 32 interactions for NEAT1 and MALAT1 networks, respectively. Almost, all the interactions in these two independent networks were supported by text-mining. Afterwards, the factors interacted with each of the two lncRNAs and were combined to find the common interacting factors (Figure 4). We identified two lncRNAs namely MIAT and TUG1 as common interacting factors while no common proteins were identified. A total of 35 factors were identified to have 75 interaction networks. Subsequently, we analyzed the expression levels of these 35 factors in ZIKV infected cells. This approach yielded a total of 11 factors with differential expression, which included 6 proteins (ABCB1, HCN2, KIAA1147, PRRG1, PSPC1, SFPQ) and 5 lncRNAs (LINC00657, MALAT1, NEAT1, SNHG11, GAS5) (Supplementary Table 3). Based on the evidence provided by the interaction networks, 3 (SFPQ, NONO, PSPC1) of 35 factors are known to be involved in common cellular and biological processes. Based on this study, 9 of 11 differentially expressed genes identified to constitute novel RNA-Protein interaction (RPI) networks (ABCB1, GAS5, HCN2, KIAA1147, LINC00657, MALAT1, NEAT 1, PRRG1, and SNHG11).

**Figure 4.**
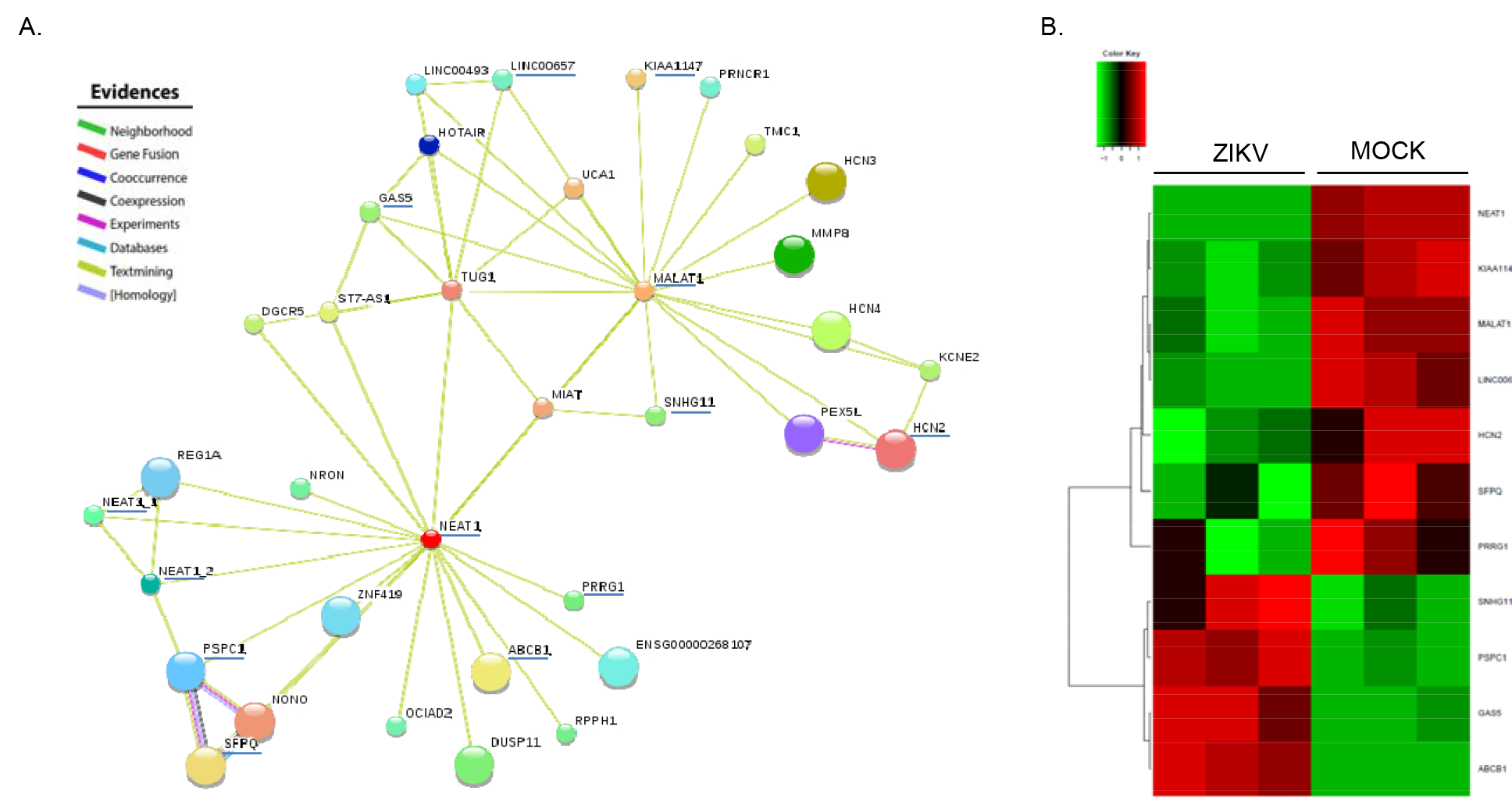
LncRNAs NEAT1 and MALAT1 interaction networks. (A) The network represents lncRNAs NEAT1 and MALAT1 interactions with 17 lncRNAs and 18 proteins, in which 5 lncRNAs and 6 proteins were co-regulated in ZIKV infected cells (underlined genes). There are 35 high confidence nodes involved in 75 interactions (PPI enrichment p-value of 0.0 and clustering coefficient of 0.856). (B) Heat map of differentially expressed genes in ZIKV infected cells, which are present in the interaction network. (Red: up-regulated genes; Green: down-regulated genes).

<u>*Comprehensive IncRNA-Protein interaction network*</u>. Proteins collectively contribute to a shared and specific function. For instance, if a single edge appears with different lines/colors between two nodes, this clearly indicates that the interaction between two given genes/nodes is supported by more than one verification [25]. It is ambiguous if current co-regulated gene data can be correlated to the predicted human lncRNA-protein networks even though it is widely known that interacting factors are frequently co-expressed [26]. With the evidence from the interacting network, we identified known co-expressing genes that are differentially regulated during ZIKV infection from our genome-wide gene expression data sets. We extended our analysis using the 11 interacting factors defined in our previous analysis for further comprehensive interacting network analysis in order to identify more high confidence coexpressing interacting factors and their possible functional associations. Bioinformatics analysis of the interacting data uncovered a network of 289 interactions between 71 factors, including 11 seeds with high statistical significance (PPI enrichment *p*-value 0; clustering co-efficient 0.699). Among the 71 interacting factors in the network, 51 factors (46 protein-coding, 2 ncRNAs and 3 lncRNAs) were differentially expressed in our study (Supplementary Table 3). As evidenced by the interaction network, 28 of 71 factors were known to be co-expressed in many cellular and biological processes. Notably, 29 of the 51 differentially expressed factors identified during ZIKV infection are novel co-expressing factors (AVL9, CYP3A5, DENND6A, ETS1, FBXO15, FOXO3, GAS5, HAX1, HCN1, HCN2, HIF1A, HSF1, HSPB1, KIAA1147, LANCL2, LINC00657, MALAT1, MAPK8, NEAT1, PIM1, PRRG1, PTBP1, RNF43, SNHG11, SP1, TAF12, TAF7, TP53, UGCG) (Supplementary Table 3). It is possible that 5 lncRNAs having interactions with 24 proteins may form post-transcriptional regulatory networks [27–29], and control fundamental cellular and developmental processes. We also observed that apart from the text-mining, the majority of the lncRNA-protein interactions were supported by experimental evidence.

<u>*Pathway analysis of interacting factors in the network*</u>. Subsequently, we performed the KEGG functional enrichment analysis for all 71 interacting factors to identify their functional associations. We noted that the factors interacting in the network were involved in 33 functional KEGG pathways, in which 79% (N=26) of the pathways consist of less than six proteins (Fig. 5; Supplementary Table 4). Interestingly, several of the enriched pathways are associated with viral infections. Furthermore, in the context of ZIKV and microcephaly, factors involved in the Neurotrophin signaling pathway (FOXO3, TP53, ABL1, JUN, MAPK8) and MAPK signaling pathway (HSPB1, TP53, FOS, JUN, MAPK8) were identified. As a result of this study, we generated a comprehensive lncRNA-protein interactions map based on the differentially expressed genes during ZIKV infection.

**Figure 5.**
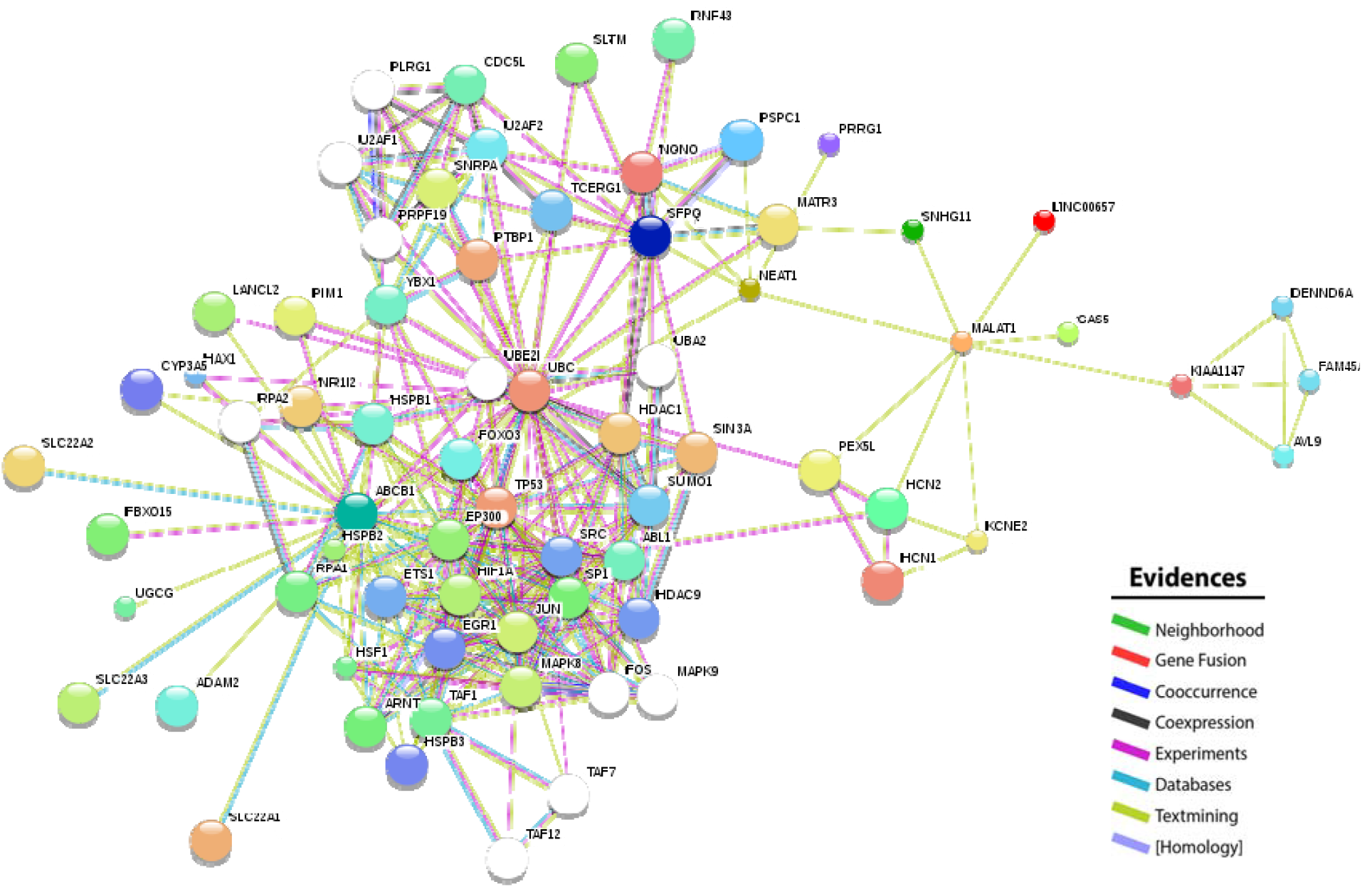
A comprehensive RNA-protein interaction network for 11 factors co-regulated in the differentially expressed gene data set. The network represents 11 factors (5 lncRNAs and 6 proteins), and additional 40 co-regulated proteins identified in our differentially expressed gene data sets. A total of 71 high confidence nodes are involved in 289 interactions, which was supported by the PPI enrichment *p*-value of 0.0 and clustering coefficient of 0.699.

MALAT1 and NEAT1 have been shown to be involved in cell proliferation and cell cycle control [30, 31]. Down-regulation of MALAT1 and NEAT1 by Zika virus can inhibit cell cycle progression. We found that the cell cycle pathways and various cell cycle genes including CDK1, Cyclin B, E2F, and p21 were deregulated in ZIKV infected cells (Supplementary Fig. 3). Our data suggests that the lncRNAs MALAT1 and NEAT1 can play important roles in the pathogenesis of Zika virus-mediated developmental diseases.

## DISCUSSION

Here, we have provided the first comprehensive analysis of the effect of ZIKV infection on cellular lncRNA gene expression. We utilized an unbiased RNA sequencing approach to profile differentially regulated genes during ZIKV infection of mammalian cells. We identified a total of 121 lncRNA genes that were differentially regulated during infection. Though over 25,000 lncRNAs are catalogued in the human genome, the functions of only a few hundred of them are known [32]. We identified that MALAT1 and NEAT1, two lncRNAs with known functions, were down-regulated upon ZIKV infection.

MALAT1 is localized at nuclear speckles. These sub-nuclear structures are involved in RNA polymerase II transcription, splicing of pre-mRNA and export of processed mRNA ([33, 34]). NEAT1 is present in the nuclear paraspeckle structures [35, 36]. Both of these lncRNAs have been shown to be upregulated in cancer cells and to be involved in cell proliferation [31, 33, 37–40]. Depletion of these lncRNAs can cause loss of cell proliferation and cell cycle arrest. ZIKV has been shown to cause cell cycle inhibition and neuronal cell death [41]. These observations allow us to formulate a hypothesis that ZIKV may regulate MALAT1 and NEAT1 for cell cycle control towards completing virus replication. This investigation was carried out in a ZIKV infected model cell line. Profiling the deregulated lncRNAs during ZIKV infection of human neuro-and retinal progenitor cells (the cell types frequently affected by ZIKV *in utero*) can yield a tissue type-specific lncRNA expression profile and functional relevance.

The majority of important molecular functions in cells are governed by interactions among proteins and other biomolecules [42, 43]. The RNA-protein interactions, which are known to facilitate post-transcriptional regulatory networks, are beginning to be understood [27–29]. Through detailed bioinformatics analysis of our differentially expressed gene database, we have identified 11 proteins/lncRNAs that form the core factors in the RPI network involving MALAT1 and NEAT1. Further comprehensive analysis revealed the formation of a larger interaction network comprised of 71 factors. This includes 51 DE genes from ZIKV infected cells. This analysis provides a comprehensive view on the complexity of lncRNA and ZIKV interaction. Although recent studies have found evidence of many RNA binding proteins and RPIs in cells [44–46], huge numbers of lncRNAs and proteins that participate in RPIs need to be characterized. Apart from regulating gene expression at the post-transcriptional level, RPIs can control many fundamental biological processes including, but not limited to, DNA replication, transcription, pathogen resistance and viral replication [47–50]. As ZIKV causes developmental abnormalities, a further detailed investigation on the functions of lncRNAs in fetal development and Zika viral disease pathogenesis could yield additional clues to prevent the teratogenic effect.

## Acknowledgments

We are thankful to Dr. Aaron Brault and Dr. Brandy Russell of the Centers for Disease Control and Prevention (CDC), USA for sharing Zika viral strain PRVABC59. We also thank Nikhil Chakravarty of Cedars-Sinai Medical Center for editing this manuscript. This work was supported by Cedars-Sinai Medical Center’s Institutional Research Award to V.A.

## Supplementary Figures and Tables

**Supplementary Figure 1.**
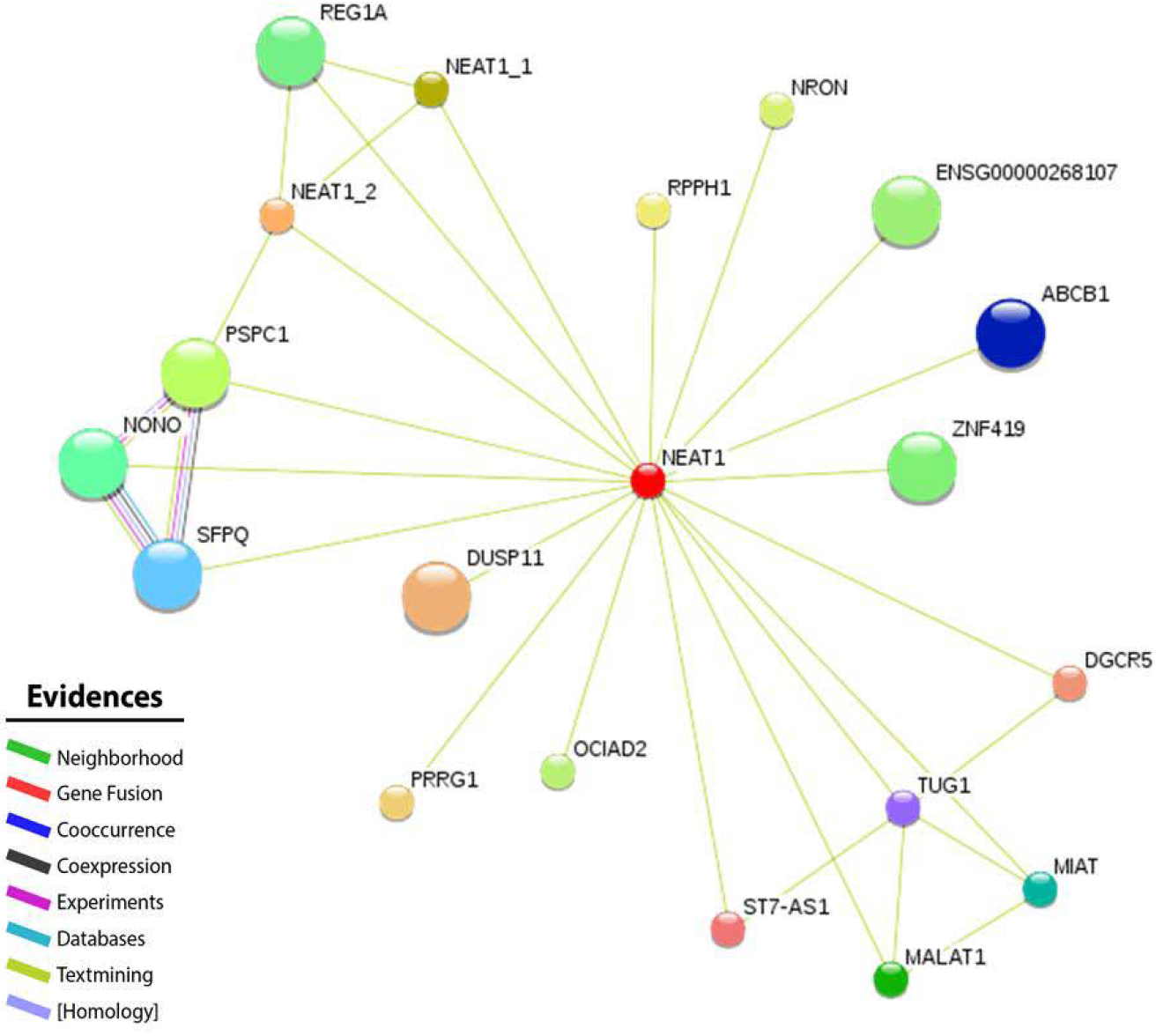
LncRNA NEAT1 interaction network. Lncrna NEAT1 interacts with 10 lncRNAs and 10 proteins, in which 2 lncRNAs and 4 proteins, were co-regulated in our data sets. A total of 20 high confidence nodes are involved in 31 interactions. This was supported by the PPI enrichment *p*-value of 0.007 and clustering coefficient of 0.895.

**Supplementary Figure 2.**
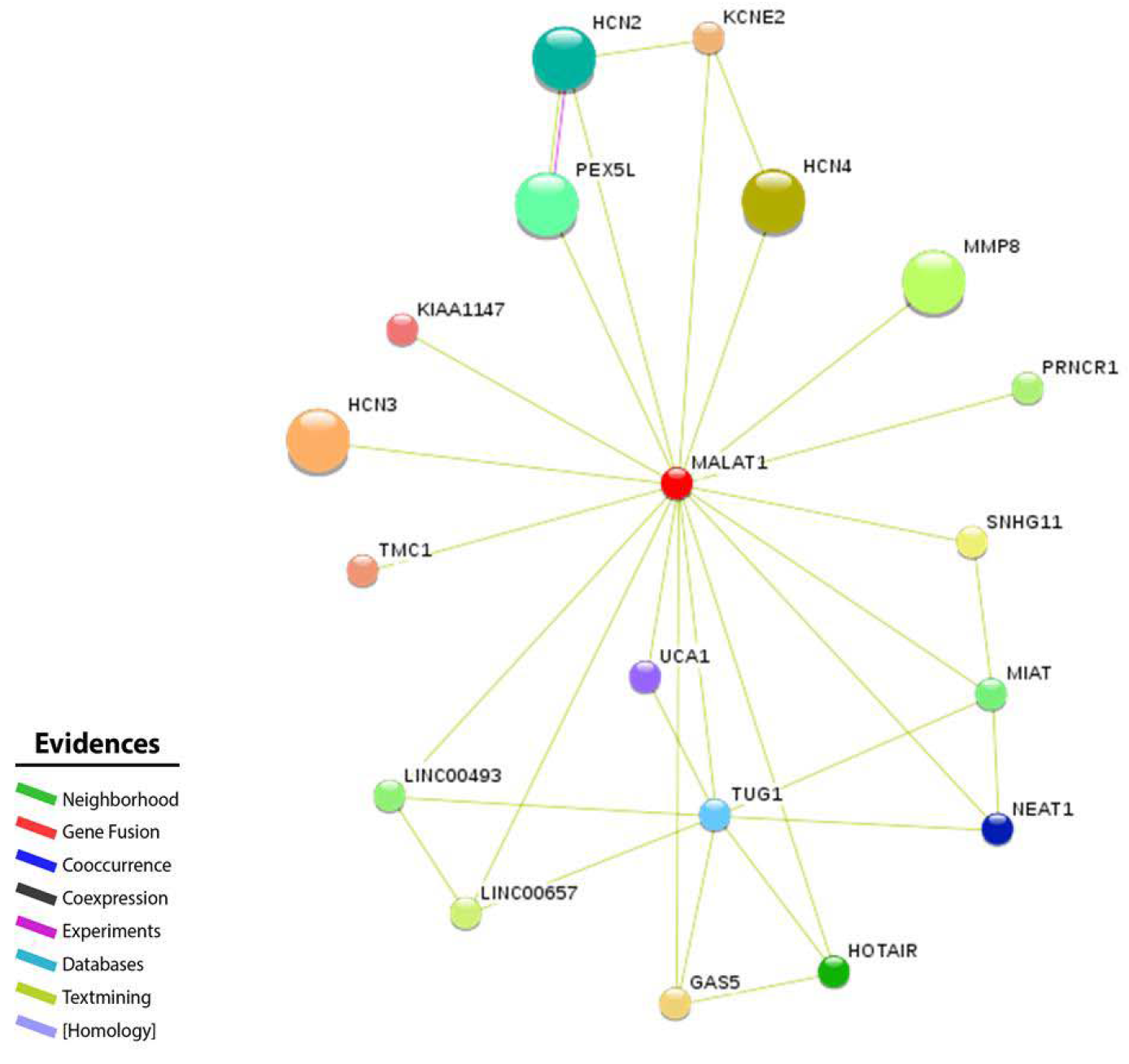
LncRNA MALAT1 interaction network. This network represents lncRNA MALAT1 interaction with biomolecules, including 11 lncRNAs and 8 proteins, in which 5 and 2 lncRNAs and proteins, respectively were co-regulated in our differentially expressed gene data sets. A total of 19 high confidence nodes are involved in 32 interactions, which was supported by the PPI enrichment *p*-value of 0.002 and clustering coefficient of 0.866.

**Supplementary Figure 3.**
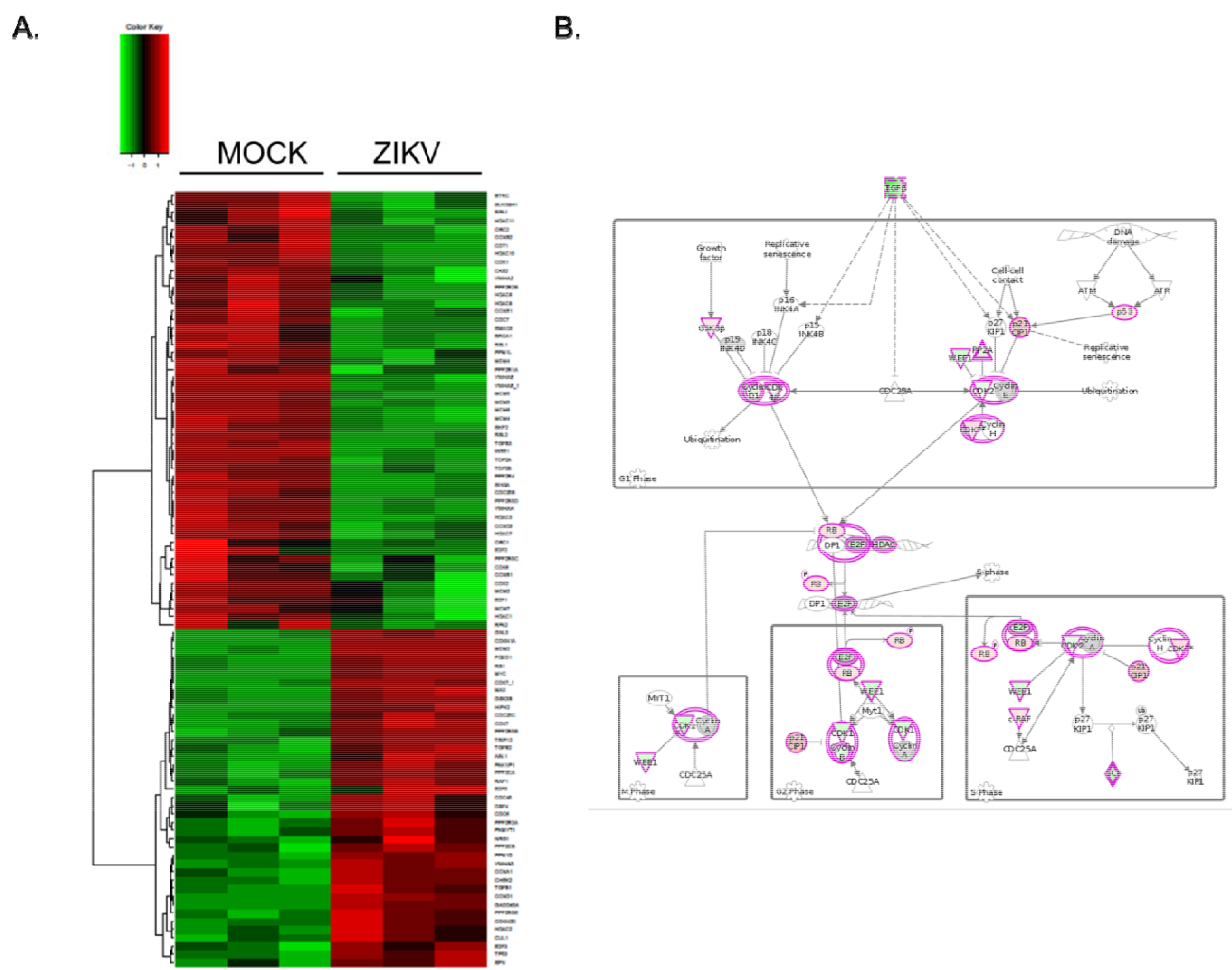
Differentially expressed (DE) genes involved in cell cycle during ZIKV infection at 48 hpi. (A) Heat map of DE genes is shown. (B) Pathway analysis of genes involved in cell cycle regulation. The positive cell cycle regulators, CDK1 and Cyclin B, are down-regulated in ZIKV infected cells.

**Supplementary Table 1.**
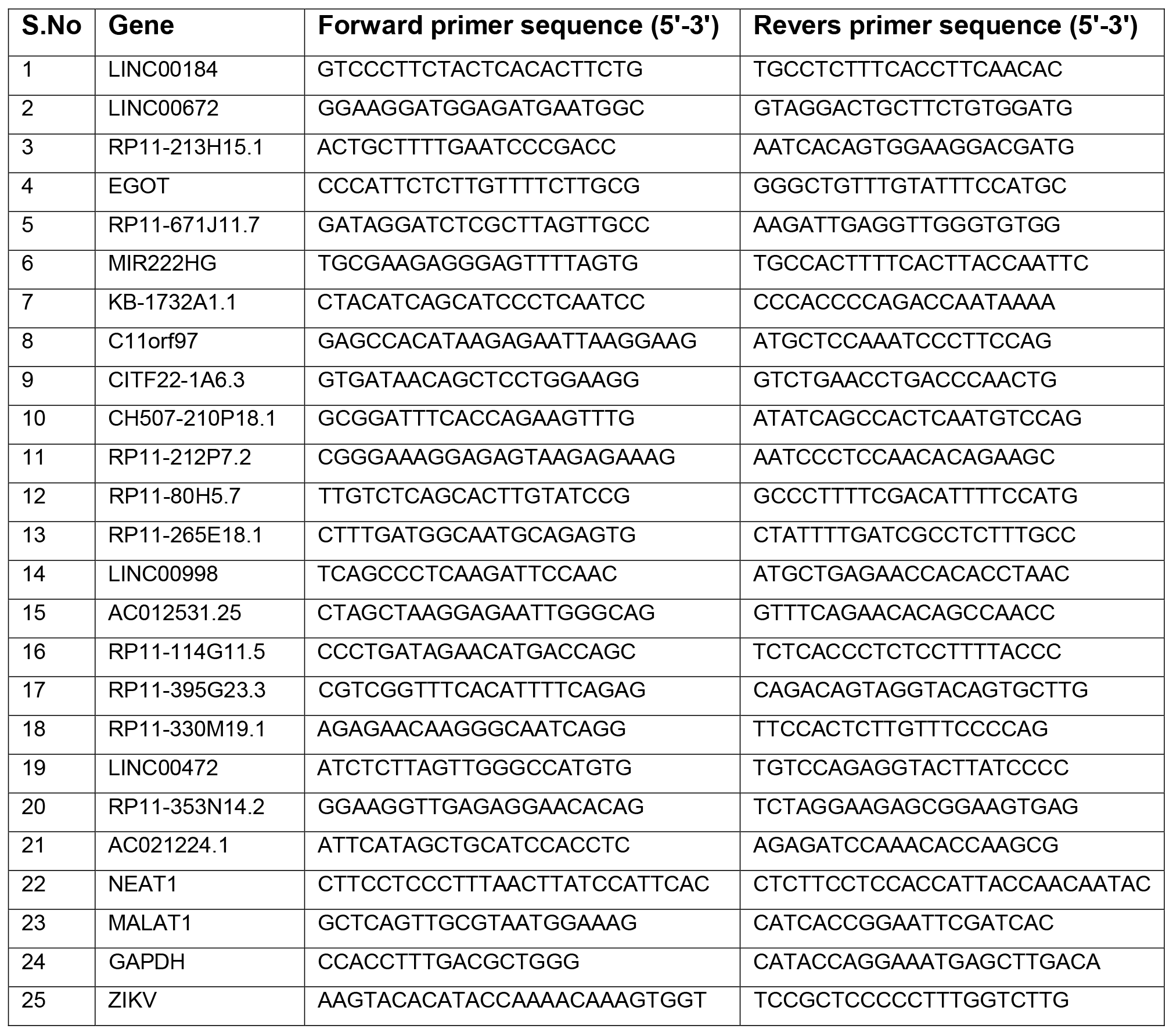
List of primers used in this study.

**Supplementary Table 2**. List of differentially expressed lncRNA genes in Zika virus infected cells at 48 hpi.

**Supplementary Table 3. Co-expressing biomolecules from the comprehensive interaction networks that were identified in our data sets**. **(A)** List of co-expressing differentially expressed 46 protein coding genes and 5 lncRNAs identified in our data sets. **(B)** List of 20 protein-coding genes that were not present in our data sets.

**Supplementary Table 4. The KEGG pathway enrichment analysis for 71 factors**. The 71 factors include 5 lncRNAs and 66 proteins from the comprehensive interaction networks (shown in Figure 5), found to be involved in 33 functional pathways.

## REFERENCES

1. Brasil, P., et al., Zika Virus Infection in Pregnant Women in Rio de Janeiro – Preliminary Report. N Engl J Med, 2016.

2. Schuler-Faccini, L., et al., Possible Association Between Zika Virus Infection and Microcephaly – Brazil, 2015. MMWR Morb Mortal Wkly Rep, 2016. 65(3): p. 59–62.

3. Mlakar, J., et al., Zika Virus Associated with Microcephaly. N Engl J Med, 2016.

4. Dick, G.W., Zika virus. II. Pathogenicity and physical properties. Trans R Soc Trop Med Hyg, 1952. 46(5): p. 521–34.

5. Dick, G.W., S.F. Kitchen, and A.J. Haddow, Zika virus. I. Isolations and serological specificity. Trans R Soc Trop Med Hyg, 1952. 46(5): p. 509–20.

6. Hamel, R., et al., Biology of Zika Virus Infection in Human Skin Cells. J Virol, 2015. 89(17): p. 8880–96.

7. Lanciotti, R.S., et al., Genetic and serologic properties of Zika virus associated with an epidemic, Yap State, Micronesia, 2007. Emerg Infect Dis, 2008. 14(8): p. 1232–9.

8. Djebali, S., et al., Landscape of transcription in human cells. Nature, 2012. 489(7414): p. 101–8.

9. Huarte, M., et al., A large intergenic noncoding RNA induced by p53 mediates global gene repression in the p53 response. Cell, 2010. 142(3): p. 409–19.

10. Kung, J.T., D. Colognori, and J.T. Lee, Long noncoding RNAs: past, present, and future. Genetics, 2013. 193(3): p. 651–69.

11. Guttman, M., et al., Ab initio reconstruction of cell type-specific transcriptomes in mouse reveals the conserved multi-exonic structure of lincRNAs. Nat Biotechnol, 2010. 28(5): p. 503–10.

12. Cabili, M.N., et al., Integrative annotation of human large intergenic noncoding RNAs reveals global properties and specific subclasses. Genes Dev, 2011. 25(18): p. 1915–27.

13. Consortium, E.P., et al., Identification and analysis of functional elements in 1% of the human genome by the ENCODE pilot project. Nature, 2007. 447(7146): p. 799–816.

14. Dinger, M.E., et al., Long noncoding RNAs in mouse embryonic stem cell pluripotency and differentiation. Genome Res, 2008. 18(9): p. 1433–45.

15. Guttman, M., et al., Chromatin signature reveals over a thousand highly conserved large non-coding RNAs in mammals. Nature, 2009. 458(7235): p. 223–7.

16. Loewer, S., et al., Large intergenic non-coding RNA-RoR modulates reprogramming of human induced pluripotent stem cells. Nat Genet, 2010. 42(12): p. 1113–7.

17. Hung, T., et al., Extensive and coordinated transcription of noncoding RNAs within cell-cycle promoters. Nat Genet, 2011. 43(7): p. 621–9.

18. Rinn, J.L. and H.Y. Chang, Genome regulation by long noncoding RNAs. Annu Rev Biochem, 2012. 81: p. 145–66.

19. Bawa, P., et al., Integrative Analysis of Normal Long Intergenic Non-Coding RNAs in Prostate Cancer. PloS One, 2015. 10(5): p. e0122143.

20. Lanciotti RS, L.A., Holodniy M, Saavedra S, del Carmen Castillo Signor L, Phylogeny of Zika virus in Western Hemisphere, 2015 [letter]. 2016.

21. Contreras, D. and V. Arumugaswami, Zika Virus Infectious Cell Culture System and the In Vitro Prophylactic Effect of Interferons. Journal of Visualized Experiments, 2016(e54767).

22. Langmead, B., et al., Ultrafast and memory-efficient alignment of short DNA sequences to the human genome. Genome Biol, 2009. 10(3): p. R25.

23. Love, M.I., W. Huber, and S. Anders, Moderated estimation of fold change and dispersion for RNA-seq data with DESeq2. Genome Biol, 2014. 15(12): p. 550.

24. Benjamini, Y. and Y. Hochberg, Controlling the False Discovery Rate: A Practical and Powerful Approach to Multiple Testing. Journal of the Royal Statistical Society. Series B (Methodological), 1995. 57(1): p. 289–300.

25. Szklarczyk, D., et al., STRING v10: protein-protein interaction networks, integrated over the tree of life. Nucleic Acids Res, 2015. 43(Database issue): p. D447–52.

26. Rhodes, D.R., et al., Probabilistic model of the human protein-protein interaction network. Nat Biotechnol, 2005. 23(8): p. 951–9.

27. Mittal, N., et al., Dissecting the expression dynamics of RNA-binding proteins in posttranscriptional regulatory networks. Proc Natl Acad Sci U S A, 2009. 106(48): p. 20300–5.

28. Kishore, S., S. Luber, and M. Zavolan, Deciphering the role of RNA-binding proteins in the post-transcriptional control of gene expression. Brief Funct Genomics, 2010. 9(5-6): p. 391–404.

29. Hafner, M., et al., Transcriptome-wide identification of RNA-binding protein and microRNA target sites by PAR-CLIP. Cell, 2010. 141(1): p. 129–41.

30. Wang, P., et al., Long noncoding RNA NEAT1 promotes laryngeal squamous cell cancer through regulating miR-107/CDK6 pathway. J Exp Clin Cancer Res, 2016. 35: p. 22.

31. Tripathi, V., et al., Long noncoding RNA MALAT1 controls cell cycle progression by regulating the expression of oncogenic transcription factor B-MYB. PLoS Genet, 2013. 9(3): p. e1003368.

32. Amaral, P.P., et al., lncRNAdb: a reference database for long noncoding RNAs. Nucleic Acids Res, 2011. 39(Database issue): p. D146–51.

33. Bernard, D., et al., A long nuclear-retained non-coding RNA regulates synaptogenesis by modulating gene expression. EMBO J, 2010. 29(18): p. 3082–93.

34. Hutchinson, J.N., et al., A screen for nuclear transcripts identifies two linked noncoding RNAs associated with SC35 splicing domains. BMC Genomics, 2007. 8: p. 39.

35. Hirose, T., et al., NEAT1 long noncoding RNA regulates transcription via protein sequestration within subnuclear bodies. Mol Biol Cell, 2014. 25(1): p. 169–83.

36. Naganuma, T. and T. Hirose, Paraspeckle formation during the biogenesis of long noncoding RNAs. RNA Biol, 2013. 10(3): p. 456–61.

37. Zeng, C., et al., Inhibition of long non-coding RNA NEAT1 impairs myeloid differentiation in acute promyelocytic leukemia cells. BMC Cancer, 2014. 14: p. 693.

38. Guo, S., et al., Clinical implication of long non-coding RNA NEAT1 expression in hepatocellular carcinoma patients. Int J Clin Exp Pathol, 2015. 8(5): p. 5395–402.

39. Gibb, E.A., et al., Human cancer long non-coding RNA transcriptomes. PloS One, 2011. 6(10): p. e25915.

40. Standaert, L., et al., The long noncoding RNA Neat1 is required for mammary gland development and lactation. RNA, 2014. 20(12): p. 1844–9.

41. Tang, H., et al., Zika Virus Infects Human Cortical Neural Progenitors and Attenuates Their Growth. Cell Stem Cell, 2016.

42. Lees, J.G., et al., Systematic computational prediction of protein interaction networks. Phys Biol, 2011. 8(3): p. 035008.

43. Wang, T.Y., et al., A predicted protein-protein interaction network of the filamentous fungus Neurospora crassa. Mol Biosyst, 2011. 7(7): p. 2278–85.

44. Baroni, T.E., et al., Advances in RIP-chip analysis: RNA-binding protein immunoprecipitation-microarray profiling. Methods Mol Biol, 2008. 419: p. 93–108.

45. Barkan, A., Genome-wide analysis of RNA-protein interactions in plants. Methods Mol Biol, 2009. 553: p. 13–37.

46. Kim, M.Y., J. Hur, and S. Jeong, Emerging roles of RNA and RNA-binding protein network in cancer cells. BMB Rep, 2009. 42(3): p. 125–30.

47. Hogan, D.J., et al., Diverse RNA-binding proteins interact with functionally related sets of RNAs, suggesting an extensive regulatory system. PLoS Biol, 2008. 6(10): p. e255.

48. Licatalosi, D.D. and R.B. Darnell, RNA processing and its regulation: global insights into biological networks. Nat Rev Genet, 2010. 11(1): p. 75–87.

49. Sola, I., et al., RNA-RNA and RNA-protein interactions in coronavirus replication and transcription. RNA Biol, 2011. 8(2): p. 237–48.

50. Li, Z. and P.D. Nagy, Diverse roles of host RNA binding proteins in RNA virus replication. RNA Biol, 2011. 8(2): p. 305–15.

